# Unsupervized identification of prognostic copy-number alterations using segmentation and lasso regularization

**DOI:** 10.1101/2022.12.14.520497

**Authors:** Alice Cleynen, Hervé Avet-Loiseau, Jill Corre

## Abstract

Identifying copy-number alteration with prognostic impact is typically done in a supervised approach, were candidate regions are user-selected (chomosome arms, oncogenes, etc). Yet CNA events may range from whole chromosome alterations to small focal amplifications or deletions, with no available approach to combine the potential prognostic impact of different aberration ranges. We propose and compare different statistical models to integrate the effects of multi-scale CNA events by exploiting the longitudinal structure of the genome, and assume that the survival distribution follows a Cox-proportional hazard model. These methods are adaptable to any cohorts screened for CNA by genome-wide assays such as CGH-array or whole-genome sequencing technologies, and with sufficient follow-up time. We show that combining a segmentation in the survival odds strategy with a lasso-regularization selection approach provides the best results in terms of recovering the true significant CNA regions as well as predicting survival outcomes. In particular, as shown on a 551 Multiple Myeloma patient cohort, this method allows to refine previously identified regions to exhibit potential novel driver genes.

## 1 Introduction

The majority of cancer cells, regardless of the cancer type, harbor copy number abnormalities (CNA) in their kariotypes. These abnormalities range from small partial gains or losses of chromosomes to entire chromosome amplifications or deletions. Despite an apparent randomness in patient kariotypes, each cancer has its own characteristics in terms of abnormalities, with some cancers such as thyroid carcinoma producing only a small number of alterations, while others, like ovarian cancer, harbor abnormalities on almost half of the karyotype [Harbers et al., 2021]. Similarly, these apparently randomly scattered anomalies within a cancer type may in fact exhibit clear patterns, as is the case for instance in Multiple Myeloma (MM) with “hyperdiploïd MM” patients (about half of them) presenting trisomies on most odd chromosomes (3,5,7,9,11,15,17,18,19,21) [Chretien et al., 2015]. In such context, the question of the association of CNAs with patient prognosis naturally arises. In fact it has even been shown [Smith and Sheltzer, 2018] that mutations in almost all cancer driver genes contain little information on patient prognosis while CNAs in these same driver genes harbor significant prognostic power.

The question of identifying the prognostic regions, on the other hand, is seldom asked, with most studies focusing on scores such as percentage of tumor genome with alteration [Hieronymus et al., 2018, Zhang et al., 2018] a list of given regions, such as cancer genes ([Smith and Sheltzer, 2018]), or studying CNAs at low resolution (whole chromosome or arm altered) [Stopsack et al., 2019, Chretien et al., 2015]. On the contrary, identifying de novo regions with prognostic impact is not addressed. Here we focus on identifying Significant Regions (SR) defined as the shortest region that will equally impact patient prognosis no matter which proportion of this region is altered (for instance a region containing a driver gene requires its full sequence to produce functional transcripts).

Even at a single CNA scale, copy-number breakpoints will differ between patients, with some patients harboring longer gains or losses than others. To reconcile copy-number events in a large cohorts analysis, a classic strategy consists in defining regions of interest on which to test the association of CNA with survival. For instance, one may partition the genome in *N* bins of equal sizes (1*kb*, 1*Mb*, etc), and assign a CNA to a patient if at least 50% of the region is gained or lost. This results in a 2 × *N* covariate vector *Xcov* where if *j* ≤ *N, Xcov*(*j*) = 1 if the patient harbors an amplification in region *j* and 0 otherwise, and if *N* ≤ *j* ≤ 2*N, Xcov*(*j*) = 1 it the patient harbors a deletion in region *j* − *N*, 0 otherwise. In most cancers, 90 % of chromosomal regions with alterations observed in more than 5% of patients are clearly unbalanced towards a large majority of gains or losses [Broët et al., 2009]. Testing for association with survival can therefore be conducted by comparing only two groups of patients, those harboring a given anomaly versus all other patients. This paper is illustrated using this approach, but all methods can easily be adapted to more than two groups.

We will use the (censoring independent) multivariate Cox proportional hazard model to describe patient’s survival as follows: for a given set of pairs (*r, e*) where *r* denotes a genomic region, and *e* a CNA event (gain or loss), for every patient *i* denote 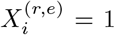 if the patient has an event *e* within region *r* and 0 otherwise. Then the hazard of patient *i* dying at time *t* is given by

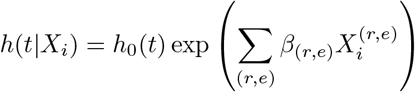

Here we review and propose statistical methods for de novo identification of (region, event) pairs with prognostic CNAs in cancer patient cohorts. The greatest strength and novelty of these methods is to produce significant survival multivariate models where the regions *r* are of variable sizes and are identified in an unsupervised framework. Resulting models may therefore combine whole chromosome arms with focal CNV events. Moreover, these methods are applicable on any CNA-assessment assays, from array comparative genomic hybridization (aCGH) to whole genome sequencing studies, and assume that patient survival times can be modeled with a multivariate Cox proportional hazard model depending only on the regions of interest, together with a given sets of covariates such as age, gender, etc.

The strategy developed is schematized in Figure 1. In a nutshell, the cohort of patient is devided into a “Selection” and a “Fit” set. The Selection set is used to aggregate initial genomic bins into meta-regions via *black box* approaches that are discussed and compared in the Results Section 2. The Fit set is then used to build a multivariate cox-proportional model using the meta-regions, and retain significant regions.

**Figure 1:**
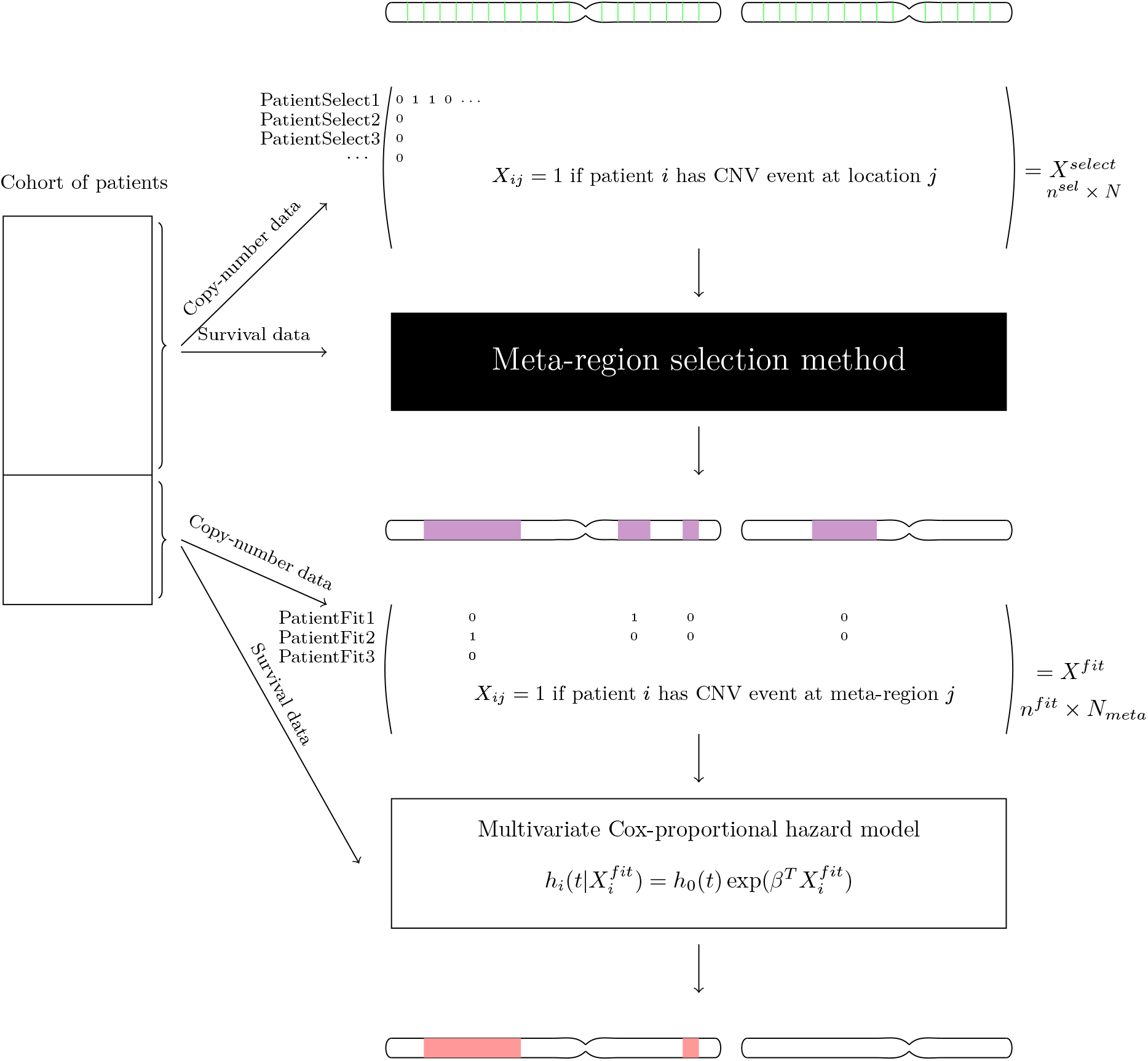
Schematized strategy. The cohort of patient is devided into a “Selection” and a “Fit” set. The Selection set is used to aggregate initial genomic bins into meta-regions via *black box* approaches that are discussed and compared in the Results Section. The Fit set is then used to build a multivariate cox-proportional model using the meta-regions, and retain significant regions.

## 2 Results

### 2.1 Black-box methods for meta-region identification

Here we supposed that the genome has been partitioned into *N* bins which we denote as {*r*_*j*_, 1 ≤ *j* ≤ *N*}, and we denote *a* an amplification event and *d* a deletion event, so that *e* ∈ {*a, d*}. For a patient *i*, we denote 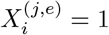 if she has an event *e* on region *r*_*j*_, and 0 otherwise.

#### Aggregation of univariate tests

A state-of-the-art strategy consists in performing univariate tests on each initial bin, and correcting for multiple testing (for instance when working at a 10kb scale, around 30 thousand regions are tested). Regions passing the significance threshold can then be aggregated into meta-regions if they are within a given distance of each-other. Here we used a Benjamini and Hochberg adjustment [Benjamini and Hochberg, 1995], which aims at controlling the False Discovery Rate (FDR), that is the expected proportion of false discoveries, and merge regions that are within 3 bins of each-other. The model is then specified as: ∀*j* ∈ {1, …, *N*}, ∀*e* ∈ {*a, d*},

- set 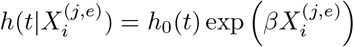 for all patients *i* in the Select group.
- test *H*_0_ : {*β* = 0} versus *H*_1_ : {*β* ≠ 0}
- if *H*_0_ is rejected, keep (*j, e*)

If (*j, e*) and (*j′*, *e*) are in the retained set and are such that |*j* − *j′*| < *s*, define 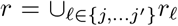 and keep (*r, e*) as a meta-region. Otherwise keep both (*j, e*) and (*j′*, *e*) separately. This method is denoted “Adj-Pval” in the simulation study.

#### Penalized multivariate Cox model

Penalized multivariate approaches try to combine all bins in a single multivariate model and penalize coefficients in order to identify relevant CNAs. The most widely spread approach is to penalize the cox coefficients by Lasso (or *L*1) variable selection, even though many other penalizations are commonly used, such as *L*2 penalization or Ridge regression. For instance, [Gui and Li, 2005] presents the LARS-Lasso procedure to identify genes whose expression are predictive of patient survival times in diffuse large B-cell lymphoma. Here again we merge significant bins (that is with non-null *β* coefficient) into a meta-region if they are less than 3 bins apart. The model is then specified as:

- set 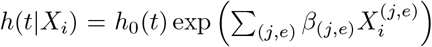 such that ∑_(*j,e*)_ |*β*_(*j,e*)_ | < *λ* for all patients *i* in the Select group.
- choose *λ* by 10-fold cross-validation
- if *β*_(*j,e*)_ ≠ 0, keep (*j, e*)

If (*j, e*) and (*j′*, *e*) are in the retained set and are such that |*j − j*^*′*^| < *s*, define 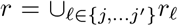 and keep (*r, e*) as a meta-region. Otherwise keep both (*j, e*) and (*j′*, *e*) separately. This method is denoted “Lasso” in the simulation study.

State-of-the-art approaches described above do not take advantage of the longitudinal structure of the data. For instance, at a high-resolution approach, it is likely that if a region is significantly associated with survival, its neighboring regions will be as well, as onco-genes may span several predefined regions, and gene families may aggregate in genomic domains. For this reason, we propose two novel approaches for the identification of copy number alterations associated with survival outcomes. Both emanate for the segmentation framework, a technique widely used for the identification of copy-number changes.

#### Segmentation of p-values

In this two-step approach we perform univariate tests on each region of interest, and post-process the p-values through a segmentation algorithm. Segmentation techniques aim at partitioning signals in homogeneous regions according to a given statistical model. In this framework, the genome will be segmented into regions of equivalent log-rank test significance. This approach will be meaningful if one expects large copy-number alterations, covering multiple successive regions of interest, that globally impact survival. This might be the case if the resolution of the study is large enough, and the disease displays large alterations, as in Multiple Myeloma.

Under the survival null hypothesis (*H*_0_ : *S*_1_(*t*) = *S*_2_(*t*)), the p-values of the log-rank test are random variables distributed under the uniform 𝒰 [0, 1] distribution. On the contrary, under the alternative hypothesis (*H*_0_ : *S*_1_(*t*) ≠ *S*_2_(*t*)), p-values will skew to the lower end of [0, 1]. While they do not allow to quantify the difference between survival distributions, similar differences will lead to similar p-values.

The proposed strategy is to apply a Gaussian segmentation model to the logit-transformed p-values as follows: ∀*j* ∈ {1, …, *N*}, ∀*e* ∈ {*a, d*},

- set 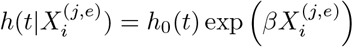 for all patients *i* in the Select group.
- test *H*_0_ : {*β* = 0} versus *H*_1_ : {*β* ≠ 0} and denote *P*_(*j,e*)_ the associated p-value
- set 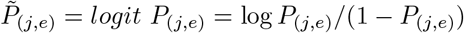

Then for *e* ∈ {*a, d*}, for *m*_*e*_ a partition of {1, …, *N*} and *r* a segment of *m*_*e*_, assume that

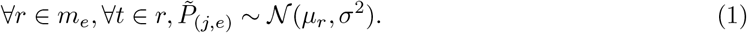

The objective is to identify the best parsimonious segmentations *m*_*a*_ and *m*_*d*_ to represent the data. Intuitively, if no CNA is significantly associated with survival, the optimal segmentation will correspond to a unique segment with mean 0 and variance *σ*^2^. Outliers corresponding to small p-values from the uniform 𝒰 [0, 1] should be identified scattered along the genome and not considered as significant chromosomal aberrations. On the contrary, significant alterations should cover successive regions of interest, and should be characterized by a segment with negative mean. We use a dynamic programming algorithm [Bellman, 1954] to recover segmentations and select the number of segments through a penalized-likelihood criteria (see Methods section). We then keep all regions with mean lower than *logit* 0.01. This method is denoted “Seg Pval” in the Simulation study

#### Segmentation of survival criteria

The log-rank test can be interpreted as a goodness of fit to the hypergeometric distribution test (see details in Appendix A.2). We propose to model instead the survival data using a Fischer non-centered hypergeometric distribution and segment the scale parameter along the genome position.

Fisher’s noncentral hypergeometric distribution is a generalization of the hypergeometric distribution where sampling probabilities are modified by weight factors. In a survival context, defining *ω* as the survival odds-ratio of groups 1 and 2, then given that each group has *m*_*k*_ individuals at risk (*k* ∈ {1, 2}) and *n* events occur at time *t*, the probability that *x* come from group 1 is given by

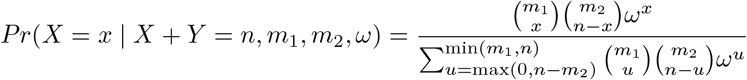

where *X* denotes the number of events occurring in group 1 and *Y* the number of events in group 2. When *ω* = 1, the Fisher’s noncentral hypergeometric distribution is the hypergeometric distribution. The log-rank test can therefore be used to test *H*_0_ : *ω* = 1 versus *H*_1_ : *ω* ≠ 1, but when *H*_0_ is rejected it gives little information on the actual odds-ratio value.

We propose a segmentation model in the odds-ratio parameter of a Fischer non-centered hypergeometric distribution to identify copy-number alterations associated with survival outcome. Using the same notations as before, the model can be written as

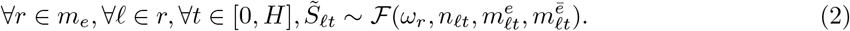

where ℱ denotes the Fisher’s noncentral hypergeometric distribution, 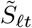 denotes the number of events observed at time *t* in the group with event *e* on bin *ℓ, n*_*ℓt*_ the number of events at time *i* observed in both groups of bin *ℓ*, and 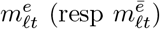 the number of patients at risk in group with event *e* (resp without event) at bin *ℓ* at time *t*.

The intuition is that neighboring regions will share the same survival odds (∼1 if the set of regions is not associated with survival) and that to infer *ω* strength can be borrowed by analyzing simultaneously segments of regions. Moreover, as opposed with segmenting p-values, the odds-segmentation directly brings information on the strength of the effect of the segments on the survival outcome. As for the previous approach, we obtain the signal segmentation using a dynamic programming algorithm.

As no theoretical guarantees exist to select the number of segments in a Fisher segmentation framework, we considered two approaches to select the final meta-regions. The first uses an arbitrary number of segments, and retains all segments with parameter *ω*_*r*_ such that |*ω*_*r*_ − 1| > 0.25. The second uses a very large number of segments and uses a penalized cox-proportional hazard model using all segments as covariates to retain the segments with non-null *β* coefficient. These two methods are denoted “Seg Odds” and “Seg+Lasso” in the Simulation study.

#### Final model for all methods

Each approach identifies a number of meta-regions of interest potentially associated with survival outcomes. We then use the “Fit” set of the cohort data to select the significant ones. For each method, we define for each patient her covariate value of meta-region (*r, e*) as 1 if she presents an event *e* covering at least a subset of meta-region *r*, and 0 otherwise. The meta-regions are then reconciled in a multivariate Cox model, and post-processed such that (i) all meta-regions with less than 1% of patients presenting a CNA are discarded, and (ii) meta-regions with p-value higher than 0.05 are successively removed starting with the highest p-value until all remaining meta-regions are significant.

### 2.2 Simulation study analysis

We propose two simulation studies (see Methods section) that differ mainly in the signal size (1000 vs 30000) which can be interpreted as the resolution at which the CNA association with survival is considered. Each simulation study is repeated 100 times for training, and tested on 100 independent datasets.

We build the long scenario in the following way. We simulate data for 500 patients and 11 significant regions (denoted SRs in what follows) associated with survival times, where we assumed the later to follow a multivariate cox-proportional model. Those SRs were fixed for all simulation datasets, with lengths varying from 15 to 750 and their strength (Cox coefficients) between −1.42 and 1. CNA events for each patient was drawn randomly in order to favor coverage of the SRs, exhibit high dependency structure, and favor coverage of additional scattered locations (see Methods section). Within each simulation dataset, coverage of the SRs varied between ∼10 and ∼180 patients. Patients were then randomly split into a Select group of 300 patients and a Fit group of 200 patients.

The short simulation study is the downscaled version of the long simulation study, where datapoints were merged into bins of size 30. Patient exhibiting a CNA on at least half a bin were counted to have a CNA on the downscaled data. Patient’s survival was not modified from the long scenario. An example of the simulation study is given in Table 1 (long scenario on the left hand-side, and its downscaled version on the right), and Supplementary Figures 1a and 2a show the number of patients with alterations along the whole signal, with the 11 SRs highlighted in blue.

**Table 1:** Long (left) simulation study parameters and its downscaled version (right)

| | Start | Length | $\beta$ | $e^\beta$ | Nb Patients | Start | Length | Nb Patients |
| --- | --- | --- | --- | --- | --- | --- | --- | --- |
| 1 | 500 | 5 | -1.42 | 0.24 | 24 | 16 | 1 | 4 |
| 2 | 3000 | 15 | -1.05 | 0.35 | 18 | 100 | 1 | 16 |
| 3 | 4250 | 750 | -0.99 | 0.37 | 98 | 141 | 26 | 101 |
| 4 | 5500 | 30 | -0.89 | 0.41 | 24 | 183 | 2 | 41 |
| 5 | 8000 | 120 | 0.42 | 1.53 | 159 | 266 | 5 | 165 |
| 6 | 10500 | 5 | 0.59 | 1.80 | 18 | 350 | 1 | 16 |
| 7 | 13000 | 15 | 0.64 | 1.89 | 153 | 433 | 1 | 133 |
| 8 | 15500 | 30 | 0.70 | 2.01 | 22 | 516 | 2 | 21 |
| 9 | 18000 | 60 | 0.72 | 2.05 | 29 | 600 | 2 | 26 |
| 10 | 20500 | 120 | 1.00 | 2.72 | 11 | 683 | 5 | 24 |
| 11 | 25500 | 15 | 1.00 | 2.72 | 18 | 850 | 1 | 17 |

Methods were evaluated with 4 criteria: an *R*2 type quantity defined as twice the difference in log-likelihood of the fitted and null cox-proportional models divided by the number of coefficients, an adjusted Rand-Index quantity (ARI) comparing the identified meta-regions and the true SRs, the SRs (at least partially) recovered by the methods, and a time-dependent AUC quantity evaluated on independent datasets and defined as in [Smith and Sheltzer, 2018] as the area under each time-dependent ROC curves where the specificity and sensitivity are

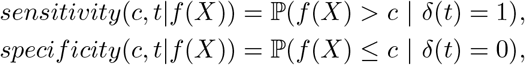

and 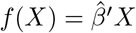 is the Cox risk score of the method, and *δ*(*t*) = 1 if an event has occurred on or before time *t. R*2, ARI, Number of coefficients and Percentage of recovered regions results are illustrated in Figures 2 and 3 for the downscaled and long scenarios respectively, while prediction scores are illustrated in Figure 4. For the long scenario, the methods requiring segmentation of the survival odds coefficient were run on only 20 simulation datasets due to computational limitations.

**Figure 2:**
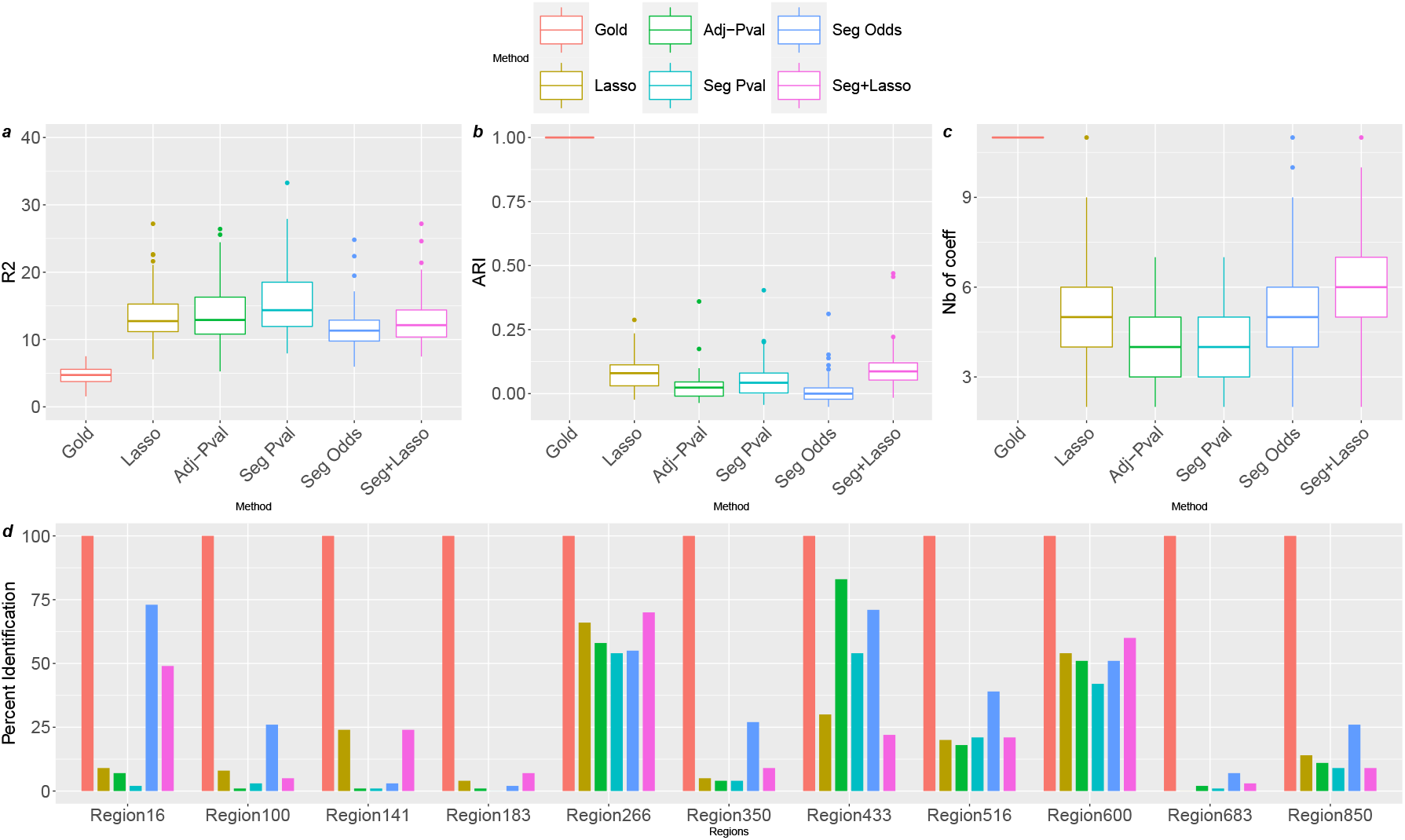
Results for 100 downscaled simulations comparing all methods. A) R2-type quantity. B) Adjusted rand-Index. C) Number of selected regions. D) Number of simulations where the regions were identified.

**Figure 3:**
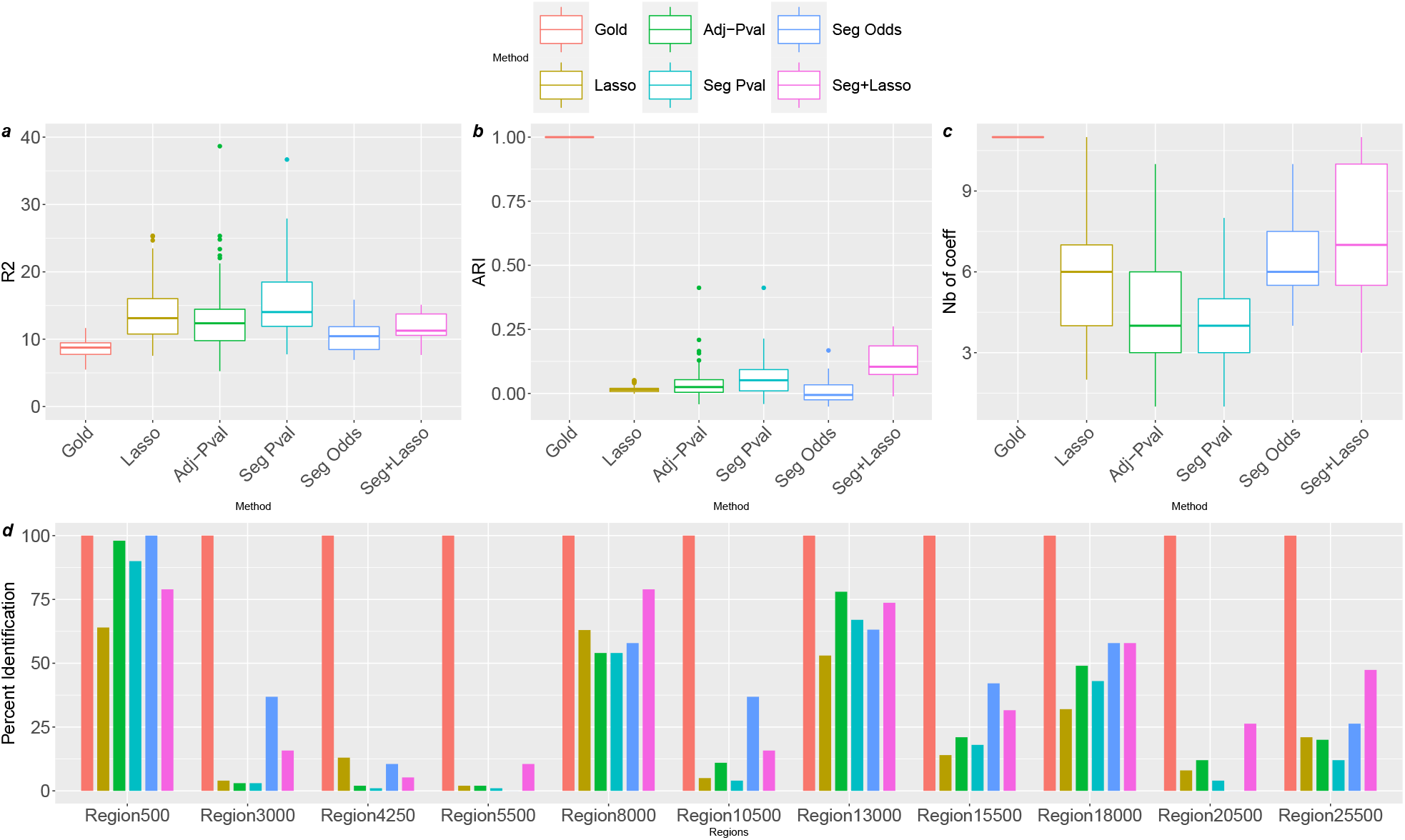
Results for 100 long simulations comparing all methods. A) R2-type quantity. B) Adjusted rand-Index. C) Number of selected regions. D) Number of simulations where the regions were identified. Methods based on the segmentation of survival odds method were only tested on 20 simulations due to computational limitations.

**Figure 4:**
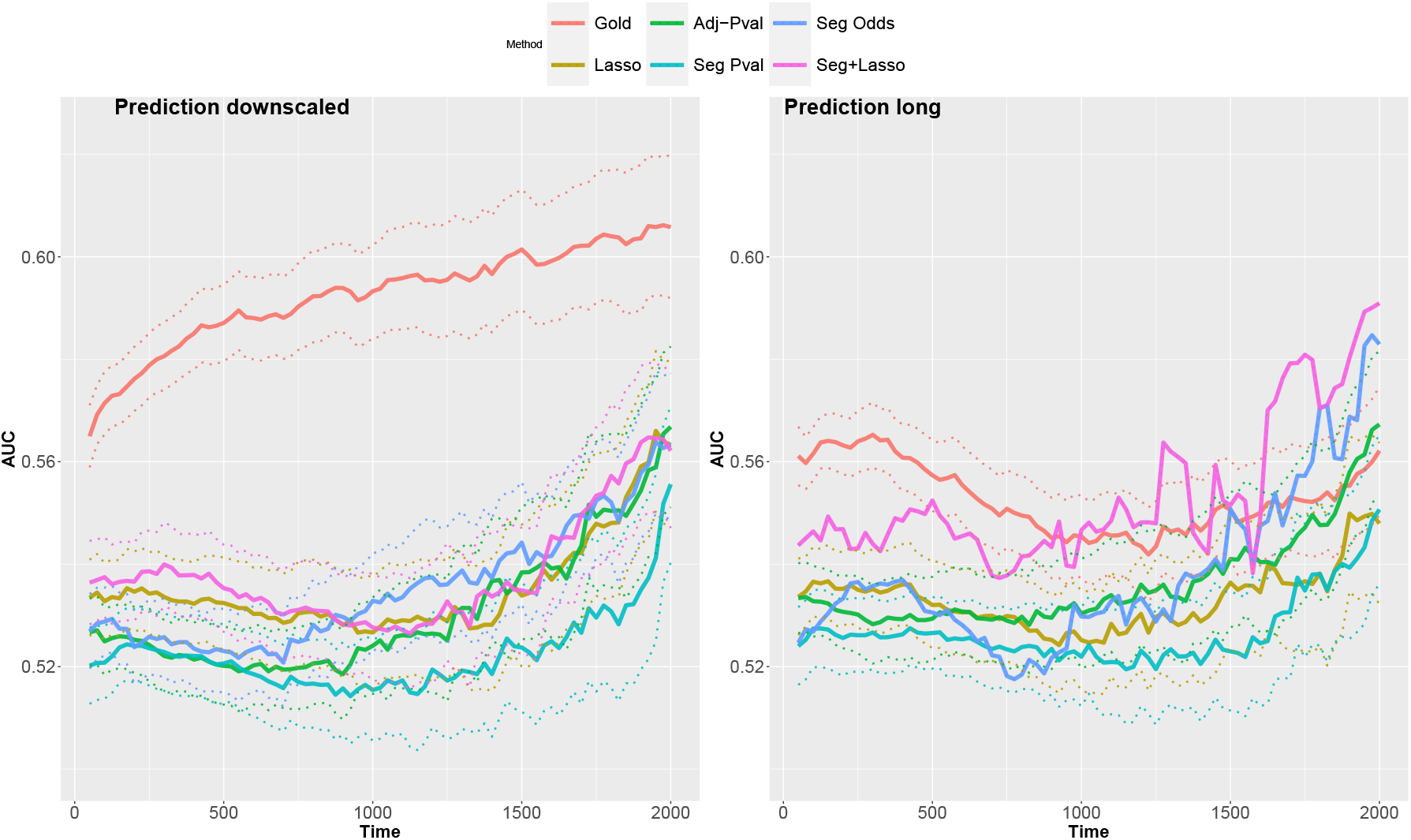
Time-dependent AUC values over the range 50 to 2000 days. Curves were averaged over 100 training simulations, each evaluated on an independent dataset. For the long scenario, methods based on the segmentation of survival odds method were only tested on 20 simulations due to computational limitations. For each method, 95% confidence intervals are given by the dotted curves.

## Results

All methods performed relatively comparably in terms of R2, ARI and number of selected coefficients, but exhibited different performances on true SR identification (see Figure 2). The methods based on p-values (multiple testing adjustment and segmentation) selected the same number of coefficients and identified the same true regions, but selecting meta-regions based on segmentation allowed to select smaller regions wich resulted in significantly better prediction scores (see Figure 4). By selecting on average one (downscaled) or two (long) more regions, the lasso approach identified significantly more true regions on the downscaled simulation scenario, but less true regions on the long simulation scenario as its inability to produce long meta-regions often resulted in selection of bins neighbours to the true SR. It’s performance in terms of ARI was therefore better for the downscaled scenario, but worse in the original simulations.

The methods based on the segmentation of the survival odds (with or without lasso post-selection) significantly identified more true SRs than all other methods. In particular, choosing meta-regions with lasso selection decreased the selected region sizes, identifying SR more precisely hence increasing the adjusted rand-index. Both methods had better prediction scores, with the combination of segmentation and lasso outbeating the gold standard in the original long scenario.

### 2.3 Identifying prognostic regions in Multiple Myeloma

As detailed in the Methods section, we applied all approaches on a cohort of 551 Multiple Myeloma patients screened with CGH arrays and with a follow-up clinical data of at least 5 years. We analyzed separately the data at the chromosome band level and at a 100kb scale, leading to signals of sizes ∼850 and ∼30000 respectively, and split the cohort in a 300 Select set and 251 Fit set. Only metaregions with at least 3 patients (about 1% of the Fit cohort) exhibiting a CNA were kept for analysis. Segmentation was run on each chromosome separately (to prevent segments from covering regions belonging to different chromosomes) but assembled before considering the multivariate cox model. We then computed the prediction scores of each method on an independent cohort of 891 Multiple Myeloma patients screened with low coverage WGS and with follow-up of 1 to 7 years (MMRF CoMMpass study). Figure 5 shows the prediction scores for the band-scale analysis.

**Figure 5:**
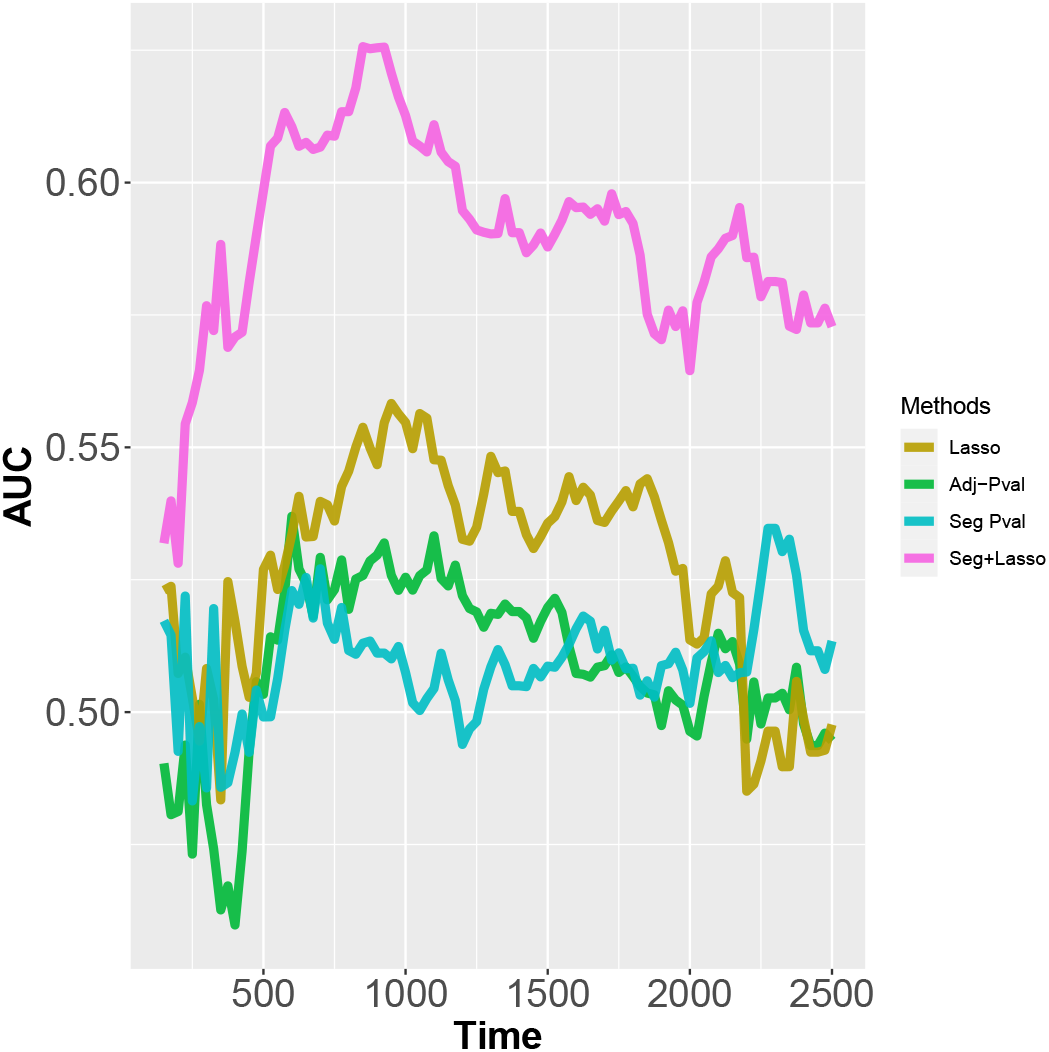
Time-dependent prediction scores of each method calibrated on a 551 MM cohort data and tested on a 891 independent MM cohort. The hybrid segmentation + Lasso approach outperforms all other methods, achieving similar AUC score as the Gold standard of the simulation study method.

Identified regions mostly agree at both scale level analysis, with the highest resolution allowing the identification of more meta-regions. In particular, we recovered the well-known Gain 1q, Deletion 1p, and Deletion 17p, and identified novel impact regions such as focal losses on chromosomes 6q or 7p. Using the segmentation-lasso hybrid approach, we were able to refine the precise locations of the impact regions to 3.4 Mb for Gain 1q, 450 Mb for Deletion 1p, or 8Mb for Deletion 17p.

## 3 Discussion

In this study we have investigated and developed methods to identify prognosis CNA events at any unsupervised scale. We have found that despite a surprising ability to identify neighboring regions of similar impact, penalized multivariate approaches fail to capture long regions, and suffer extensively from “close miss”, *i*.*e*. selecting regions very close to true prognostic regions. Methods based on correcting p-values for multiple testing, on the other hand, tend to select very large regions, failing at identifying the origin of their impact on survival. Replacing the multiple-testing correction by a segmentation framework improves significantly this behavior, at the risk of failing to identify some regions of lower impact. Using Fischer’s non-centered hypergeometric distribution to model survival within a segmentation model avoids the high dependency issue of univariate testing, identifies almost systematically all regions of interest, but suffers from the difficulty of selecting the correct number of segments. Combining this approach with a Lasso-penalization selection improves significantly the ability to recover true regions of interest (outbeating all other methods).

Surprisingly, downscaling the data to wider bins seemed to affect only the performances of methods using lasso regularization. As expected, the standard lasso approach performed poorly on the long datasets, failing to identify neighboring bins as a single meta-region. However, combining a segmentation approach to create initial meta-regions of similar prognosis impact and a lasso-regularization scheme to select the significant ones provided the best results, especially in the smallest scale situation. Put together, these approaches benefit from the precision of the small scale without suffering from the inability to merge neighboring bins.

Interestingly, in the simulation study, the regions that were least often identified (uniformly by all methods) were in fact those with the highest impact (*β* = −1.42 for the first region, and *β* = 1 for the tenth region), also corresponding to regions with low number of patients. The longest region was almost systematically at least partially identified by all methods, while, as expected, the lasso approaches almost systematically failed to identify the shortest regions, most often selecting instead regions close by. Increasing the cohort size to 1000 patients slightly increased the coverage of the identified SR, but did not provide significant improvements (see Supplementary Materials). Finally, the predictive power of all methods, including the Gold approach (true model) was very poor, with AUC curves averaging at 0.5, the equivalent of a “flip a coin” strategy in balanced scenarios (note that here, especially at early time-points, the number of patients with events is drastically lower than those without event). Here again, a similar simulation study with no censoring survival times did not improve the predictive power of the methods (see Supplementary Materials). These extensive simulation studies suggest that our results on the Multiple Myeloma cohorts are as good as one could hope for, with the Segmentation-lasso approach reaching prediction scores as good as the Gold model in the simulation study.

## 4 Materials and Methods

### 4.1 Data sources

The dataset considered in this study is based on a homogeneous series of 551 patients newly diagnosed with multiple myeloma. This study was approved by the Centre de Recherche en Cancérologie de Toulouse ethic committee. DNA was extracted from frozen sections using the Nucleon DNA extraction kit (BACC2, Amersham Biosciences, Buckinghamshire, UK), according to the manufacturer’s procedures. For each tumor, two micrograms of tumor and reference genomic DNAs were directly labeled with Cy3-dCTP or Cy5-dCTP respectively and hybridized onto aCGH containing 32,000 DOP-PCR amplified overlapping BAC genomic clones (average size of 200 kb) providing tiling coverage of the human genome. Hybridizations were performed using a MAUI hybridization station, and after washing, the slides were scanned on a GenePix 4000B scanner. For this analysis, we only considered BAC genomic clones mapping to automosomal chromosomes. The aCGH signal intensities were normalized using a two-channel microarray normalization procedure. CNV calling was done using the `segmentCGH` function of the `rCGH R` package with default parameters (minimum segment size 10kb and smoothing of raw signal). Two analysis scales were selected, one at the chromosome band level, the other every 100kb. For each patient, a region was called as gain (respectively loss) if at least 1*/*2 of the region was amplified (resp lost) for the chromosome band scale, and at least 4*/*5 of the region for the 100kb scale. This created two signals of length 855 for the chromosome band scale, and another two of length 30970 for the 100kb scale.

### 4.2 Penalized Cox proportional hazard models

Lasso-penalized multivariate Cox models were fit using the glmnet function of the `glmnet R` package with family = “cox” and “alpha”=1. The penalization parameter *λ* was chosen via cross-validation using the default cv.fit function corresponding to a ten-fold cross validation. We then considered two choices of *λ* corresponding to the minimum cross-validated error (denoted *smin* in our analyses) and to the largest value of lambda such that the error is within one standard error of the minimum (denoted 1*se* in our analyses). As results with the 1*se* selection criteria were consistently poorer than results with the *smin* criteria, the former was dropped in all analyses.

### 4.3 Univariate Cox proportional hazard models

For each region of interest, patients were divided into two groups depending on whether they exhibited a CNA intersecting the region or not (for the Multiple Myeloma cohort, gains and losses were treated separately). Cox-propotional models were fit using the `coxph` function of the `R` package `survival` [Terry M. Therneau and Patricia M. Grambsch, 2000]. For the Multiple-testing approach, we used the Benjamini and Hochberg procedure [Benjamini and Hochberg, 1995] implemented in the `p.adjust` function of the `stat R` package. A threshold of 0.05 was used to select regions post-correction, and regions less than 3 (respectively 10 for the high resolution analyses) points apart were merged together to form a meta-region.

### 4.4 Segmentation of p-values

For the segmentation of p-values approach, P-values from the univariate cox proportional hazard tests were further logit transformed using the *logit*(*x*) = log(*x/*1 − *x*) function. No selection was performed. A homoscedastic Gaussian segmentation model was then fit using the `R` package `Segmentor3IsBack` [Cleynen et al., 2014]. The number of segments was selected using a Birgé-Massart penalization criteria [Birgé and Massart, 1997] developed for Gaussian changepoint models [Lebarbier, 2005]. All segments with parameters smaller than −4.59 = *logit*(0.01) were kept for further analysis (multivariate cox-proportional model). Theoretical indications on the validity of segmenting p-values are given in Appendix B.

### 4.5 Segmentation of survival odds

The segmentation model developed for the survival odds is based on the following pseudo-likelihood (see Appendix C for its origin):

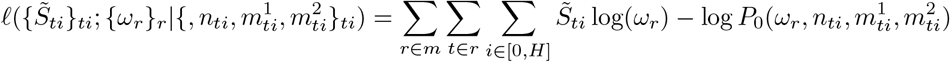

where 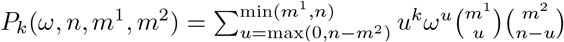.

As there is no closed form for the maximum-likelihood estimator of *ω*, and no explicit update equation of *ℓ* when extending segments, running a gradient descent algorithm within a segmentation algorithms would be computationally too expensive. Instead, we maximize *ω* over a large fixed grid within a dynamic programming algorithm. As there are no Birgé-Massart selection criterion for Fisher’s hypergeometric distribution, we selected the number of segments using a modified-BIC criterion [Zhang and Siegmund, 2007]. As this criterion tended to select very few segments (8 on average) we instead selected a fixed number of segments 30 in all simulation studies, and 8 per chromosome in our MM application. More insight is given in Appendix C, and all codes are available at https://github.com/acleynen/Segmentation-Lasso-Survival.

### 4.6 Simulation Study

The high-resolution simulation study was constructed first, and further downscaled to the lower-resolution simulation study. We thus describe here the strategy for building the long simulation study.

To simulate patient CNAs we considered 3 types of regions:

- 4 long CNAs (average size 1500) frequently shared by half patients. Three SRs were included in those long CNAs : the longest one (mimicking a full chromosome arm) and two short ones.
- Frequent short CNAs (18 regions with average size 15) that can be shared between patients. These regions covered all SRs but the longest one (hence including the two short SRs also covered by the long CNAs).
- Random short CNAs randomly (average size 30) distributed along the signal.

Patients were divided in two groups. The first half exhibited CNA in a subset (number drawn from a uniform (1, 4) distribution) of the long CNAs, as well as 0 or 1 (ℬ (1*/*2) probability) of the frequent short CNA, and a number of random CNAs drawn from a Poisson distribution with mean 1. The second half did not exhibit any long CNAs, but *n*_*f*_ ∼ *𝒫* (2) frequent short CNAs, and *n*_*r*_ ∼ *𝒫* (2) random short CNAs. Length and starting sites of all CNAs were randomly chosen by drawing size proportions in uniforms 𝒰 ([0.5, 1]) and start sites uniformly around the 4 long CNAs and frequent short CNAs definition starting sites.

To simulate patient survival, we considered the multivariate cox-proportional model where all coefficients of non significant regions were set to 0, hence for each patient *i*,

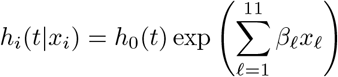

where *ℓ* ∈ {1, … 12} denotes a significant region, and *x*_*ℓ*_ = 1 if the patient has a CNA covering part of region *ℓ*, and zero otherwise. The *β*_*ℓ*_ where randomly drawn from a mixture of 𝒩 (1, 0.5) and 𝒩 (−1, 0.5) distributions. The baseline function *h*_0_ was chosen as a Weibull distribution. Finally we introduced censoring times with an exponential distribution so that the censoring rate was around 15%.

### 4.7 Evaluation criteria

Final multivariate Cox survival models were computed using the coxph function from the `survival R` package, and the *R*2 quantities were computed using the log-likelihood output of this function. *ARI* quantities were evaluated using the `adj.rand.index` function from the `pdfCluster R` package, and AUC values were computed using the auc function from the `pROC R` package.

## A Survival models and tests

### A.1 Reminder on the Cox proportional model

We consider the usual survival model where to each patient is associated a set (*y*_*i*_, *x*_*i*_, *δ*_*i*_) where *y*_*i*_ is the survival time, *x*_*i*_ is the set of covariates (for instance amplification or deletion of a region of the genome) and *δ*_*i*_ is the censoring variable, equal to 1 if the event is observed, or to 0 in the case of right-censoring. We let *t*_1_ < *t*_2_ < … < *t*_*m*_ be the increasing list of unique failure times, and *j*(*i*) denote the index of the patient whose event occurs at time *t*_*i*_. The Cox model ([Cox, 1972]) assumes that

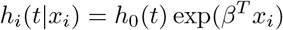

where *h*_*i*_(*t*|*x*) is the hazard of patient *i* at time *t* given his set of covariates *x*_*i*_ and *h*_0_(*t*) is a baseline function (common to every patient).

Inference on *β* is obtained by maximization of the partial likelihood

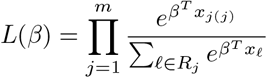

where *R*_*j*_ denotes the set of patients still at risk at time *t*_*j*_, *i*.*e*. those for which *y*_*i*_≥ *t*_*j*_.

The coefficients of *β* can then be interpreted as the instant relative risk associated to the covariates.

#### Inference

Rewriting the partial likelihood as

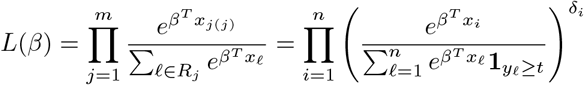

then the log partial likelihood becomes

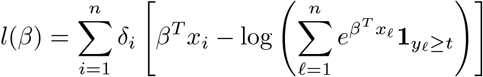

We can define the score function *U* (*β*) = (∂*l/*∂*β*_1_, …, ∂*l/*∂*β*_*p*_) with the equation

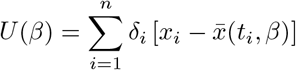

where

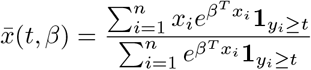

is the weighted average of the covariate vectors of individuals at risk at time *t*. The information matrix is defined by

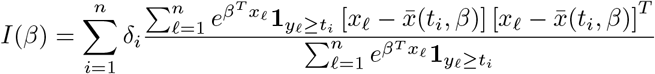

Solving *U* (*β*) = 0 can be obtained using Newton-Raphson algorithm, and standard asymptotic normal distribution is obtained for the maximum likelihood estimate

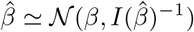

#### Test

When establishing a Cox model, one is often interested in comparing the survival distributions of two groups of individuals differing by their value in a particular covariate (for instance copy-number gain in one group at one location compared to no alteration in the other group). To simplify the problem, one may consider that the covariate value *x*_(*j*)_ is null in the reference (no alteration) group and is equal to 1 in the other group. In this case, the Cox model assumes *S*_1_(*t*) = *S*_0_(*t*) and *S*_2_(*t*) = *S*_0_(*t*) exp(*β*_(*j*)_).

The test hypothesis *S*_1_(*t*) = *S*_2_(*t*) is therefore equivalent to testing *β*_(*j*)_ = 0. Dropping the index (*j*) for sake of simplicity, one obtains

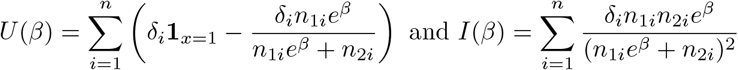

In particular in *β* = 0, one gets

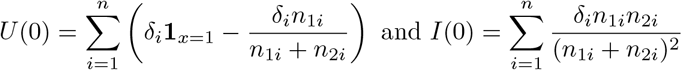

A test statistic is therefore obtained as 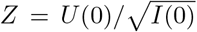 which is approxim<u>ately</u> standard normal when *n* tends to infinity. In fact one can also show that 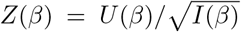 is asymptotically standard normal when *β* is the true parameter value. The interested reader will find more details in [Lawless, 2011].

### A.2 Reminder on the log-rank test

Log-rank test are widely used to compare the lifetime distributions of two groups of individuals, for instance when only one covariate separates patients in two groups. It tests the equality of the survival functions: *H*_0_ : *S*_1_(*t*) = *S*_2_(*t*) under the hypothesis of proportional hazards. This test is very popular as it requires no hypothesis on the distribution of the lifetimes, and is computed very easily: let *N*_1_ and *N*_2_ denote the number of individuals in groups 1 and 2 respectively, *n*_1*i*_ (resp *n*_2*i*_) denote the number of individuals at risk in group 1 (resp 2) at time *t*_*i*_, and *O*_1*i*_ (resp *O*_2*i*_) denote the observed number of events in group 1 (resp 2) at time *t*_*i*_. Then under the null hypothesis the statistic

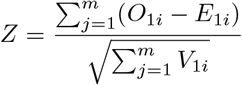

converges to that of a standard normal distribution as *m* tends to infinity, where

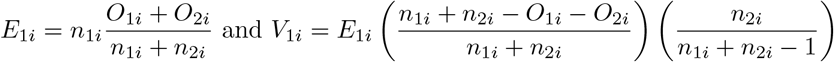

#### Log-rank test and Cox proportional hazards

The log-rank test is a simple way of testing equality of the cox-proportional survival distributions of two groups of individuals differing by only one covariate. One can first notice that, using the notations above, 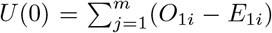. Then under the assumption of a discrete-time model allowing for ties in the failure times, the Information *I*(0) can be written as 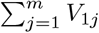 (see [Lawless, 2011]). Hence in practice, discretizing the time of any given data-set, the log-rank test is equivalent to testing the null cox-proportionnal hazard model.

#### Log-rank test and hypergeometric distribution

The null hypothesis of a log-rank test is basically assuming that at each event-time *t*_*i*_, the probability that the event occurs in one given group is purely dictated by the size of each group. In other words, at time *t*_*i*_, if *A*_1_ denotes the number of events occurring in group 1, then *A*_1_ follows a hypergeometric distribution with parameters (*n*_*i*1_ + *n*_*n*2_, *n*_*i*1_, *O*_*i*1_ + *O*_*i*2_). Then the *Z*^2^ statistic is Pearson’s chi-square test of the contingency table of the *A*_1*t*_ and *A*_2*t*_.

## B Theoretical validity of segmenting p-values

The idea of partitioning p-value signals into homogeneous regions has previously been explored using Hidden Markov Models, for instance in the context of differential gene expression testing [Sun and Tony Cai, 2009], or using post-hoc FDR control in a spatial multiple-testing setting [Sun et al., 2015]. The former approach describes a two-state HMM framework, where one states represent the null hypothesis (for instance no association between a region and survival in our scenario) and the other any alternative (region positively or negatively affecting survival). While the provide technical guarantees on the control of the FDR, their approach requires being able to compute the forward-backward density variables, which is technically intricate in our scenario.

In the second work, the authors also provide FDR control in a Bayesian computational framework, where the procedure starts with a point-wise test statistic, combines regions of interest based on the p-values, and extend an MCMC scheme to estimate an oracle quantity *T*_*OR*_ that controls the FDR.

Here we consider a simple segmentation model to partition the signal based on the logit transformed p-values. Assuming that neighboring regions sharing an impact on survival should have a similar survival distribution, their associated test should have close p-values. Segmentation models have shown very powerful to detect regions of homogeneous signals, as for instance to detect copy number alterations [Zhang and Hao, 2015, Ruan et al., 2019], as their inference do not require independence of the data within segments, only between segments. This implies that the regions of interest might be chosen very small even if it leads to identical log-rank test performed (as typically the same set of patients will constitute the groups tested for survival).

Noting that under the null hypothesis the p-values are distributed along a Uniform distribution on [0, 1], the logit-transformed p-values will approximately be distributed according to a centered Gaussian distribution with variance 3. On the other hand, genome locations within a significant region should correspond to identical survival distributions *H*_1_, hence lead to identical p-values when testing for *H*_0_. The randomness of patients CNA breakpoints introduces randomness in the *H*_1_ distribution as some patients will switch group within the SI. The logit-transformed p-values can therefore be considered as distributed from a Gaussian random variable with mean specific to the SI, and variance that will depend on the study resolution. We choose to segment the data with a homoscedastic Gaussian model. In practice, this variance may be estimated from the data using the median of estimates from sliding windows.

## C Theoretical insights on segmenting survival odds

Recall that we model 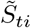, the number of events occurring at time *i* in region *t* in group 1, as following a Fisher noncentral hypergeometric distribution. In the segmentation context of model (2) and assuming groups at each region *t* are independent, the complete log-likelihood can be written as

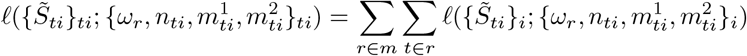

As the number of patients at risk at time *i* + 1 depend on the events at time *i*, for a given *t* the 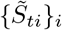 are not independent, hence the likelihood of model (2) is intractable. Instead we propose to consider the conditional log-likelihood

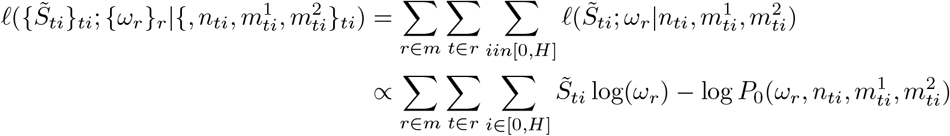

where 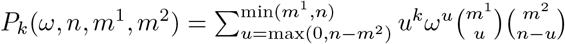

Here again this conditional log-likelihood cannot be easily optimized as there is no closed form for the maximum likelihood estimator of *ω*_*r*_ and gradient descent algorithm are difficult to implement within segmentation algorithms. [McCullagh and Nelder, 2019] propose a moment-like estimator for *ω* as

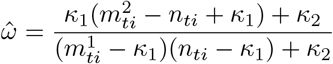

where 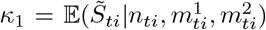 and 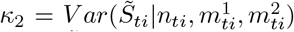. Our strategy consists in finding the optimal segmentation with respect to 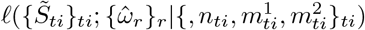. For sake of computational complexity, we choose to compute the quantities

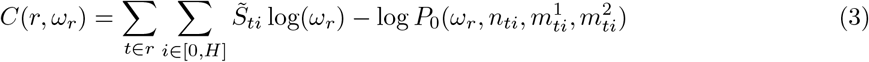

for a large grid 𝒢_*ω*_ of *ω*_*r*_ values, then compute the 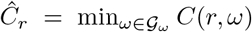 and find the optimal segmentation via dynamic programming based on the *Ĉ*_*r*_ quantities. This strategy can be efficiently implemented, since the quantities *C*(*r, ω*) are simple sums of the point quantities 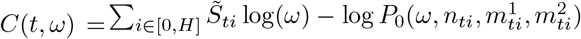 for points falling within segment *r*. Hence the main computational bottleneck of our approach is the computation of the *C*(*t, ω*). For a grid of 214 *ω* values, the full segmentation strategy takes on average 30 minutes for a signal of size 1000, and 3 days for a signal of size 30000 on a regular laptop using one CPU.

## References

Bellman, R. (1954). The theory of dynamic programming. Bulletin of the American Mathematical Society, 60(6):503–515.

Benjamini, Y. and Hochberg, Y. (1995). Controlling the false discovery rate: a practical and powerful approach to multiple testing. Journal of the Royal statistical society: series B (Methodological), 57(1):289–300.

Birgé, L. and Massart, P. (1997). From model selection to adaptive estimation. In Festschrift for lucien le cam, pages 55–87. Springer.

Blanchard, G., Neuvial, P., and Roquain, E. (2020). Post hoc confidence bounds on false positives using reference families. The Annals of Statistics, 48(3):1281–1303.

Broët, P., Tan, P., Alifano, M., Camilleri-Broët, S., and Richardson, S. (2009). Finding exclusively deleted or amplified genomic areas in lung adenocarcinomas using a novel chromosomal pattern analysis. BMC Medical Genomics, 2(1):1–11.

Chretien, M.-L., Corre, J., Lauwers-Cances, V., Magrangeas, F., Cleynen, A., Yon, E., Hulin, C., Leleu, X., Orsini-Piocelle, F., Blade, J.-S., et al. (2015). Understanding the role of hyperdiploidy in myeloma prognosis: which trisomies really matter? Blood, The Journal of the American Society of Hematology, 126(25):2713–2719.

Cleynen, A., Koskas, M., Lebarbier, E., Rigaill, G., and Robin, S. (2014). Segmentor3isback: an r package for the fast and exact segmentation of seq-data. Algorithms for Molecular Biology, 9(1):1–11.

Cox, D. R. (1972). Regression models and life-tables. Journal of the Royal Statistical Society: Series B (Methodological), 34(2):187–202.

Grinde, K. E., Arbet, J., Green, A., O’Connell, M., Valcarcel, A., Westra, J., and Tintle, N. (2017). Illustrating, quantifying, and correcting for bias in post-hoc analysis of gene-based rare variant tests of association. Frontiers in genetics, 8:117.

Gui, J. and Li, H. (2005). Penalized cox regression analysis in the high-dimensional and low-sample size settings, with applications to microarray gene expression data. Bioinformatics, 21(13):3001–3008.

Harbers, L., Agostini, F., Nicos, M., Poddighe, D., Bienko, M., and Crosetto, N. (2021). Somatic copy number alterations in human cancers: An analysis of publicly available data from the cancer genome atlas. Frontiers in oncology, page 2877.

Hieronymus, H., Murali, R., Tin, A., Yadav, K., Abida, W., Moller, H., Berney, D., Scher, H., Carver, B., Scardino, P., et al. (2018). Tumor copy number alteration burden is a pan-cancer prognostic factor associated with recurrence and death. Elife, 7:e37294.

Lawless, J. F. (2011). Statistical models and methods for lifetime data, volume 362. John Wiley & Sons.

Lebarbier, É. (2005). Detecting multiple change-points in the mean of gaussian process by model selection. Signal processing, 85(4):717–736.

McCullagh, P. and Nelder, J. A. (2019). Generalized linear models. Routledge.

Ruan, J., Liu, Z., Sun, M., Wang, Y., Yue, J., and Yu, G. (2019). Dbs: a fast and informative segmentation algorithm for dna copy number analysis. BMC bioinformatics, 20(1):1–14.

Smith, J. C. and Sheltzer, J. M. (2018). Systematic identification of mutations and copy number alterations associated with cancer patient prognosis. elife, 7:e39217.

Stopsack, K. H., Whittaker, C. A., Gerke, T. A., Loda, M., Kantoff, P. W., Mucci, L. A., and Amon, A. (2019). Aneuploidy drives lethal progression in prostate cancer. Proceedings of the National Academy of Sciences, 116(23):11390–11395.

Sun, W., Reich, B. J., Tony Cai, T., Guindani, M., and Schwartzman, A. (2015). False discovery control in large-scale spatial multiple testing. Journal of the Royal Statistical Society: Series B (Statistical Methodology), 77(1):59–83.

Sun, W. and Tony Cai, T. (2009). Large-scale multiple testing under dependence. Journal of the Royal Statistical Society: Series B (Statistical Methodology), 71(2):393–424.

Terry M. Therneau and Patricia M. Grambsch (2000). Modeling Survival Data: Extending the Cox Model. Springer, New York.

Zhang, L., Feizi, N., Chi, C., and Hu, P. (2018). Association analysis of somatic copy number alteration burden with breast cancer survival. Frontiers in Genetics, 9:421.

Zhang, N. R. and Siegmund, D. O. (2007). A modified bayes information criterion with applications to the analysis of comparative genomic hybridization data. Biometrics, 63(1):22–32.

Zhang, Z. and Hao, K. (2015). Saas-cnv: a joint segmentation approach on aggregated and allele specific signals for the identification of somatic copy number alterations with next-generation sequencing data. PLoS computational biology, 11(11):e1004618.

