## Supplementary figures for "Unsupervized identification of prognostic copy-number alterations using segmentation and lasso regularization"

### 1 Introduction

Figure 1: Results on a long simulation study (30000 datapoints), with true SR indicated in blue: number of individuals with CNA event A). Coefficients estimated from lasso-penalized multivariate cox model with selected meta-regions indicated by vertical bars B). Log Benjamini and Hochberg corrected p-values from univariate log-rank test and meta-regions selected between vertical bars C). Logit-transformed p-values from univariate log-rank test and meta-regions selected by segmentation algorithm between vertical bars D). Fischer non-central hypergeometric odds coefficient estimated from a segmentation algorithm, and meta-regions selected between vertical bars E). Fischer non-central hypergeometric odds coefficient estimated from a segmentation algorithm, and meta-regions selected by multivariate lasso-penalized cox model indicated by vertical bars F)

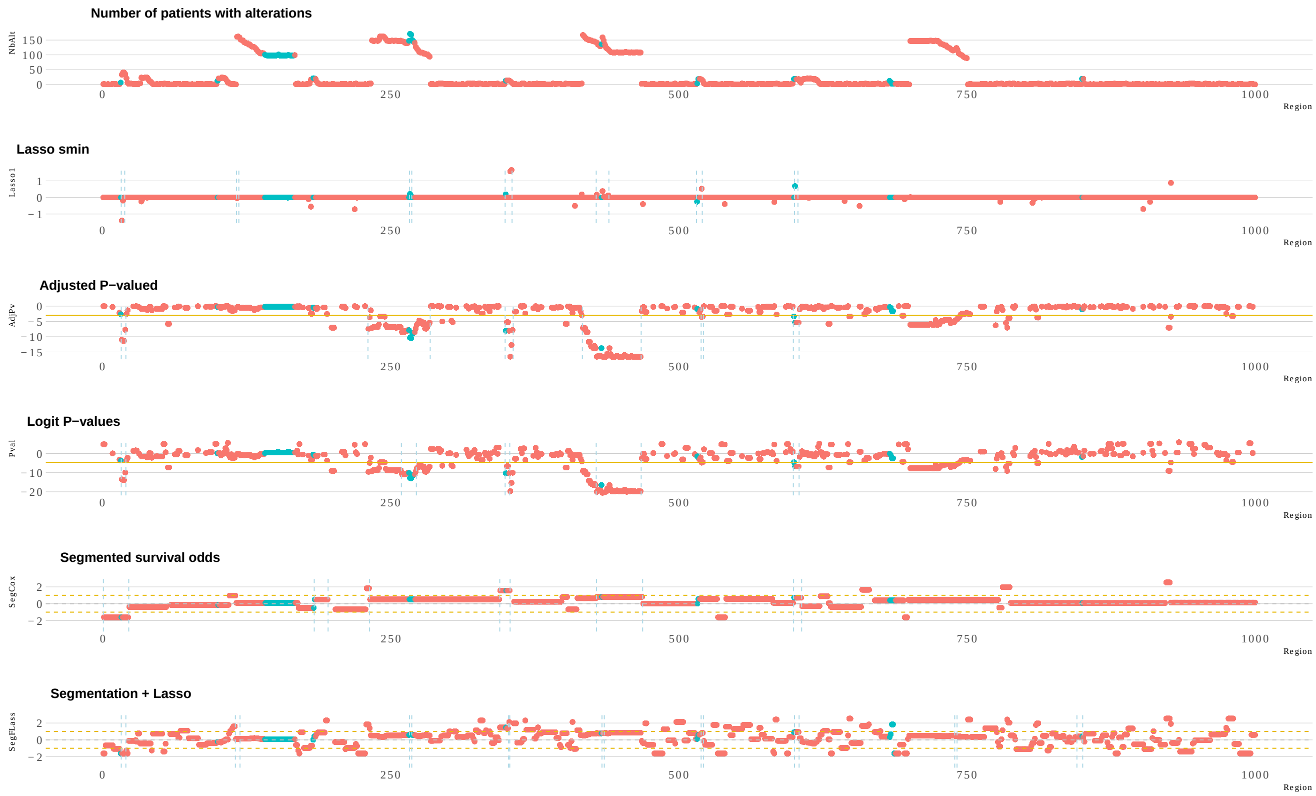


Figure 2: Results on a downscaled simulation study (1000 datapoints), with true SR indicated in blue: number of individuals with CNA event A). Coefficients estimated from lasso-penalized multivariate cox model with selected meta-regions indicated by vertical bars B). Log Benjamini and Hochberg corrected p-values from univariate log-rank test and meta-regions selected between vertical bars C). Logit-transformed p-values from univariate log-rank test and meta-regions selected by segmentation algorithm between vertical bars D). Fischer non-central hypergeometric odds coefficient estimated from a segmentation algorithm, and meta-regions selected between vertical bars E). Fischer non-central hypergeometric odds coefficient estimated from a segmentation algorithm, and meta-regions selected by multivariate lasso-penalized cox model indicated by vertical bars F)

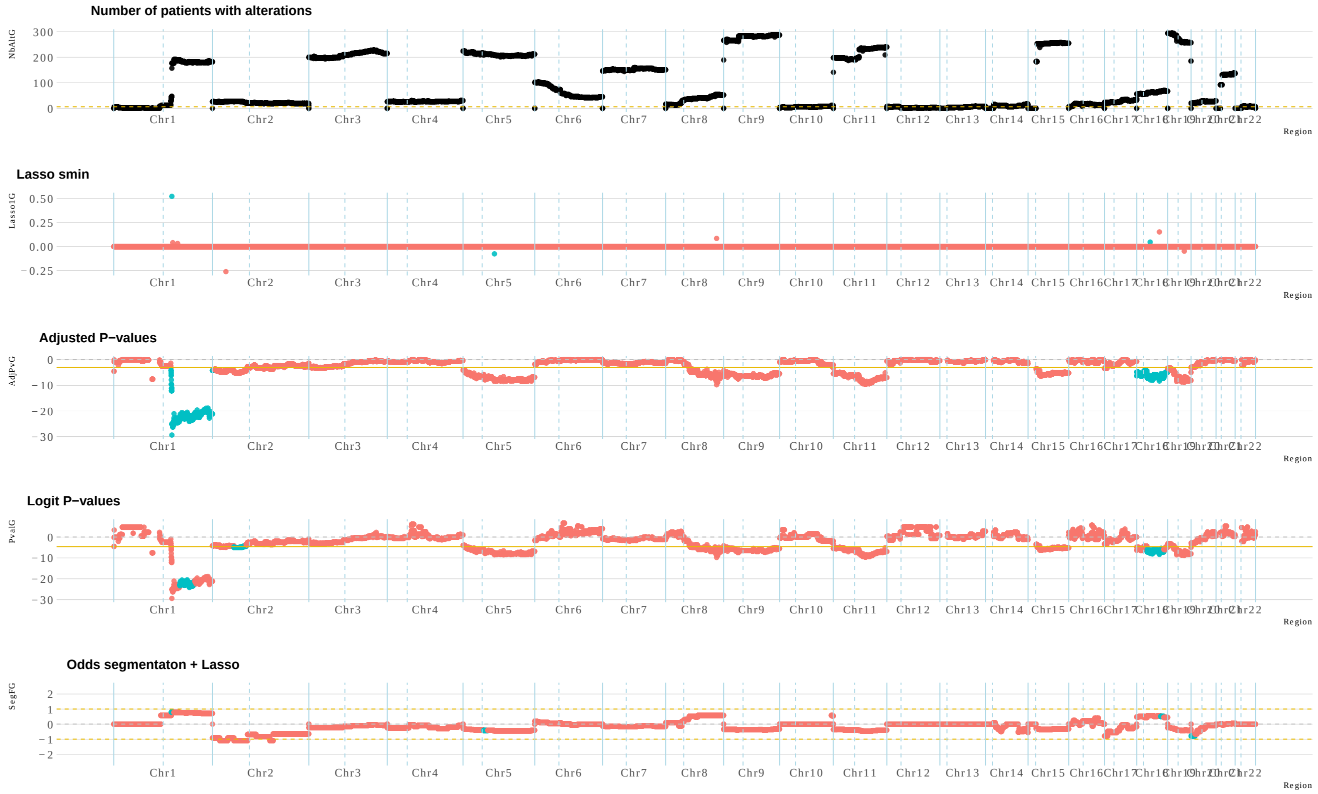


Figure 3: Results on a Multiple Myeloma cohort study of 551 patients with 10kb scale: number of individuals with amplification events event A). Coefficients estimated from lasso-penalized multivariate cox model with selected meta-regions indicated in blue B). Log Benjamini and Hochberg corrected p-values from univariate log-rank test and meta-regions selected indicated in blue C). Logit-transformed p-values from univariate log-rank test and meta-regions selected by segmentation algorithm indicated in blue D). Fischer non-central hypergeometric odds coefficient estimated from a segmentation algorithm, and meta-regions selected by multivariate lasso-penalized cox model indicated in blue E)

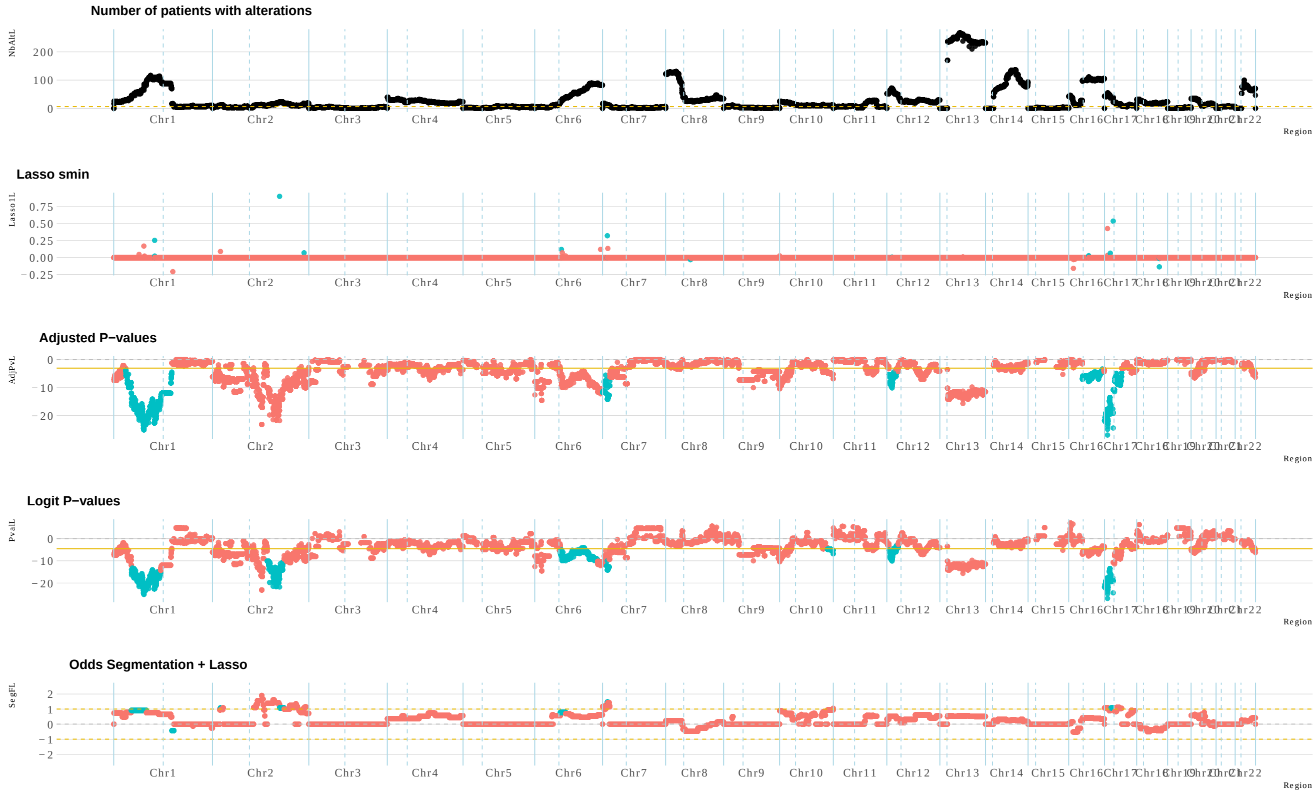


Figure 4: Results on a Multiple Myeloma cohort study of 551 patients with 10kb scale: number of individuals with deletion events event A). Coefficients estimated from lasso-penalized multivariate cox model with selected meta-regions indicated in blue B). Log Benjamini and Hochberg corrected p-values from univariate log-rank test and meta-regions selected indicated in blue C). Logit-transformed p-values from univariate log-rank test and meta-regions selected by segmentation algorithm indicated in blue D). Fischer non-central hypergeometric odds coefficient estimated from a segmentation algorithm, and meta-regions selected by multivariate lasso-penalized cox model indicated in blue E)

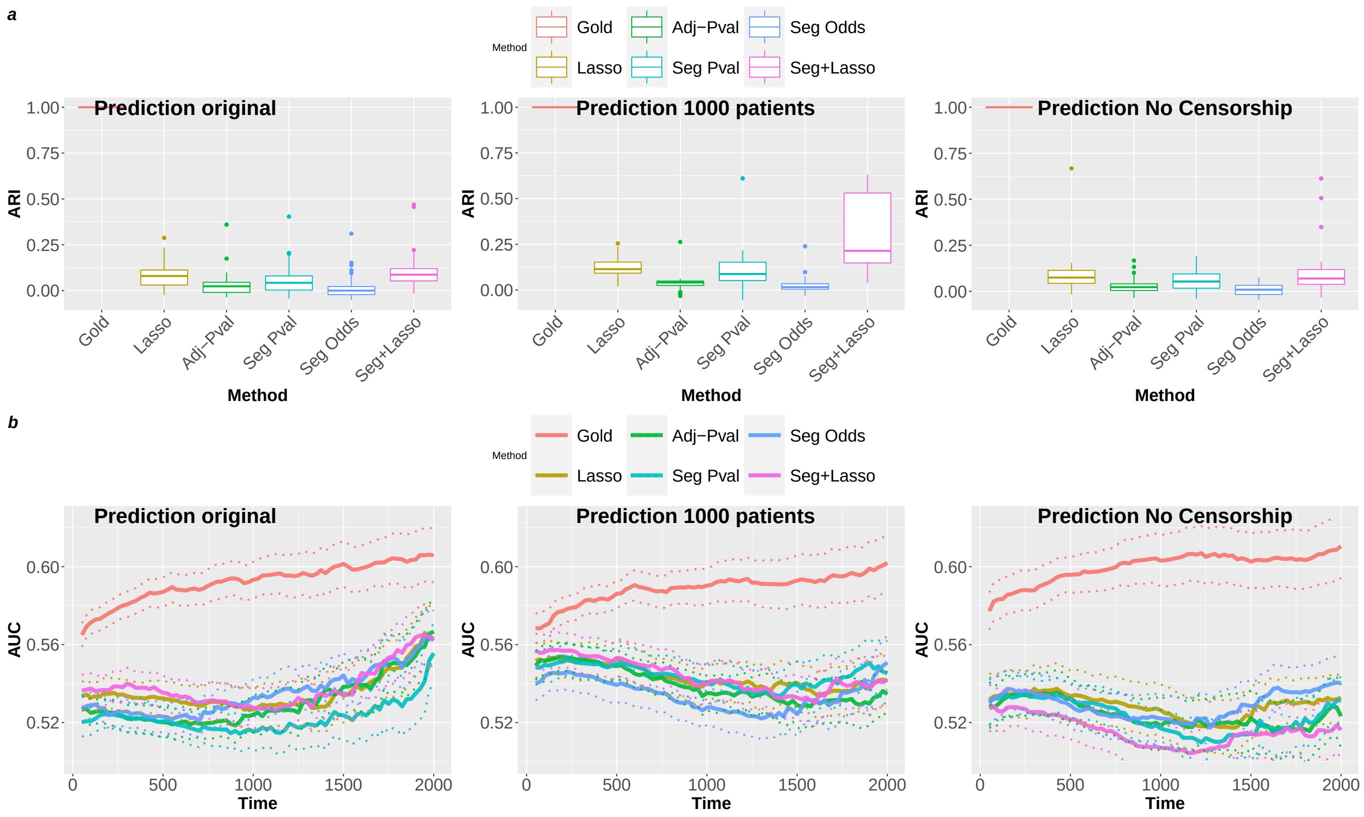


Figure 5: Adjusted Rand-Index and Prediction scores of all methods on extended simulation studies: orignial study is presented in the left column, study with 1000 simulated patients in the middle column, and study with no survival censorship in the right column.
